# *In vivo* AAV9-SB-CRISPR screen identifies fatty acid elongase ELOVL5 as a pro-resolving mediator in lung inflammation

**DOI:** 10.1101/2025.06.11.659216

**Authors:** Shuo Liu, Jiaming Wang, Liangpo Li, Jun Zhu, Xuetao Cao

**Affiliations:** Department of Immunology, Center for Immunotherapy, Institute of Basic Medicine, Peking Union Medical College, Chinese Academy of Medical Sciences, Beijing 100005, China; Institute of Immunology, College of Life Sciences, Nankai University, Tianjin 300071, China

**Keywords:** Inflammation resolution, Polyunsaturated fatty acids, ELOVL5, Lipid metabolism, Influenza, Lung inflammation

## Abstract

Timely resolution of inflammation is essential to prevent tissue damage and maintain homeostasis. Immunometabolism is critical for innate immunity and inflammation. However, how metabolic enzymes and metabolites contribute to inflammatory resolution remains largely unknown. To identify the key metabolic mediators of inflammation resolution, we generated an AAV9-Sleeping Beauty CRISPR library comprising 17090 sgRNAs targeting 2682 mouse metabolic genes. We then conducted an *in vivo* CRISPR screen in type II alveolar epithelial cells (AECIIs)-specifically expressing Cas9 mice and uncovered a very long chain fatty acid elongase, ELOVL5, that promoted the resolution of lung inflammation after influenza virus infection. Deficiency of *Elovl5* in mouse lung epithelial cells impaired lung inflammation resolution and tissue repair phenotype both *in vitro* and *in vivo*. Mechanistically, ELOVL5 bound to STING, inhibiting TBK1 interaction and translocation to the Golgi. These effects ultimately reduced STING-mediated inflammation and promoted AKT1-mediated tissue repair. In addition, ELOVL5 decreased eicosanoid levels in AECIIs to promote lung inflammation resolution. Supplement with ELOVL5 downstream products reversed the increased expression of inflammatory cytokines caused by *Elovl5* deficiency. These results support an unrevealed mechanism for polyunsaturated fatty acid metabolism in the resolution of innate inflammation and provide paths toward treating inflammatory diseases through manipulating cellular lipid metabolism.

## Introduction

Efficient termination of the innate response and resolution of inflammation are critical for tissue repair and homeostasis maintenance^1, 2^. Unraveling the intricate mechanisms underlying inflammation resolution is instructive for treating inflammatory diseases, including chronic infection, autoimmune diseases, and cancer^3^. Emerging insights into immunometabolism have advanced our understanding of the dynamic interplay between metabolic pathways and immune responses^4, 5, 6^. Indeed, immunometabolism has revealed that metabolic alterations can influence inflammation and offers a powerful tool to identify therapeutic targets and biomarkers for inflammatory diseases^7,8^. However, given complex metabolic regulation in different tissues and disease stages, how metabolic pathways coordinate to regulate the active resolution of inflammation and tissue repair remains largely unclear.

Pulmonary inflammation is a hallmark of various respiratory diseases, including coronavirus disease 2019 (COVID-19), asthma, lung cancer, etc. Controlling unresolved inflammation is critical for treating these pulmonary inflammatory diseases, which however, remains challenging due to a lack of effective and specific targets. Many cells, cytokines, lipid mediators have been found to be involved in regulating lung inflammation ^9, 10, 11^. Among them, type II alveolar epithelial cells (AECIIs) are central to promoting inflammation resolution and tissue repair. Indeed, AECIIs can produce surfactants to prevent alveolar collapse, and cytokines to influence immune responses^12^. Besides, by activating WNT/β-catenin, YAP/TAZ and other cellular signalings, AECIIs can self-renew and differentiate into AECⅠs to facilitate lung regeneration in response to injury^13, 14, 15^. Although data indicate that inflammatory signals contribute to the regeneration of alveolar epithelium^16, 17^, the endogenous mechanism that instructs the resolving response in AECIIs remains undetermined. Whether these cells undergo metabolic reprogramming during inflammation, and how they integrate metabolic and immune signals to promote inflammation resolution and timely restore the barrier integrity of the lung epithelium are unanswered questions.

Current data indicate that polyunsaturated fatty acid (PUFA) metabolism plays an important role in the resolution of inflammatory responses. PUFAs, such as omega-3 and omega-6 fatty acids, can be enzymatically converted to specialized pro-resolving mediators (SPMs) that actively promote the termination of inflammation and facilitate tissue repair^18^. However, the precise mechanisms by which PUFA metabolism orchestrates these processes remain incompletely understood. It has been reported that lipid biosynthetic genes are induced in the late program after TLR4 activation to inhibit NF-κB-mediated proinflammatory cytokine expression in macrophages^19^. In addition, agonists of liver X receptors (LXRs), which control the expression of PUFA biogenesis enzymes, promote 2’3’-cGAMP degradation and inhibit STING-mediated inflammatory signaling^20^. These studies link PUFA biogenesis to inflammation resolution and indicate relevant metabolic enzymes, such as elongases and desaturases, may also play an unexpected role in immune regulation, especially in AECII-mediated lung inflammation. Unraveling these challenges is essential for demonstrating the therapeutic potential of navigating PUFA metabolism in the control of inflammatory diseases.

In this study, we aimed to identify the essential metabolic enzymes and metabolites that regulate AECII-mediated lung inflammation resolution and tissue repair *in vivo*, and then examined the underlying mechanism for this metabolic regulation. Therefore, we used an AAV-CRISPR vector carrying the Sleeping Beauty transposon system for *in vivo* genome editing and screening. We selected 2682 mouse metabolic pathway genes from the KEGG and Recon2 databases, generated an AAV9-Metabolic CRISPR library containing 17090 sgRNAs targeting these genes, and conducted an *in vivo* CRISPR screen in a mouse model of influenza virus (IAV) infection-induced lung inflammation in AECII-specifically expressing-Cas9 mice. This screen identified fatty acid elongase 5 (ELOVL5) as an essential gene for lung inflammation resolution. ELOVL5 belongs to the very long chain fatty acid elongase family and has recently been implicated in tumor-infiltrating neutrophil reprograming^21^. However, whether and how ELOVL5 regulates the inflammation-resolving response in AECII and lung inflammation is unclear. By using AAV-LungM3-carrying sgRNA or shRNA to specifically knock down *Elovl5* in AECIIs, we validated that ELOVL5 promoted lung inflammation resolution and tissue repair in both CAG-Cas9 and C57BL/6 mice after IAV infection. Furthermore, we found that ELOVL5 promoted lung inflammation resolution as a dual immune-metabolic regulator through both interacting with STING and regulating lipid metabolism. These results uncover immune-metabolic crosstalk during pulmonary inflammation and provide foundational evidence for developing therapeutic strategies that promote inflammation resolution and enhance lung repair by targeting cellular lipid metabolism.

## Results

### Generation and validation of *in vivo* infection efficiency of AAV9-Metabolic CRISPR library

To test the function of metabolic pathway components in regulating inflammation resolution and tissue repair in the context of the tissue microenvironment, we constructed an AAV-mediated CRISPR library to edit and screen metabolic genes *in vivo*. We first generated an AAV-CRISPR vector carrying the sleeping beauty transposon system (AAV-SB) to mediate the *in vivo* integration of sgRNAs into the genome as previously reported^22, 23^ (**Fig. 1a**). We then designed a library containing 2682 mouse metabolic pathway genes from the KEGG and Recon2 databases, which included most synthetase and catabolic enzymes (**Data S1**). We designed six sgRNAs for each metabolic gene (except for *Uxs1* since only four usable sgRNAs can be designed) and 1000 nontargeting sgRNAs to generate a metabolic sgRNA library containing 17090 sgRNAs in total (**Data S1**). We pool-synthesized this sgRNA library, assembled it into an AAV-SB vector to generate the AAV-Metabolic CRISPR vector library, and verified the coverage and representation of these sgRNAs in the vector library by next-generation sequencing. The results showed that 99.99% of sgRNAs (17088) were present (**Fig. 1b**), and the skewness of this library (90% ranking/10% ranking) was 3.82 (**Fig. 1c**). These data suggested that this AAV-Metabolic CRISPR vector library had sufficient coverage and uniformity for screening.

**Figure 1.**
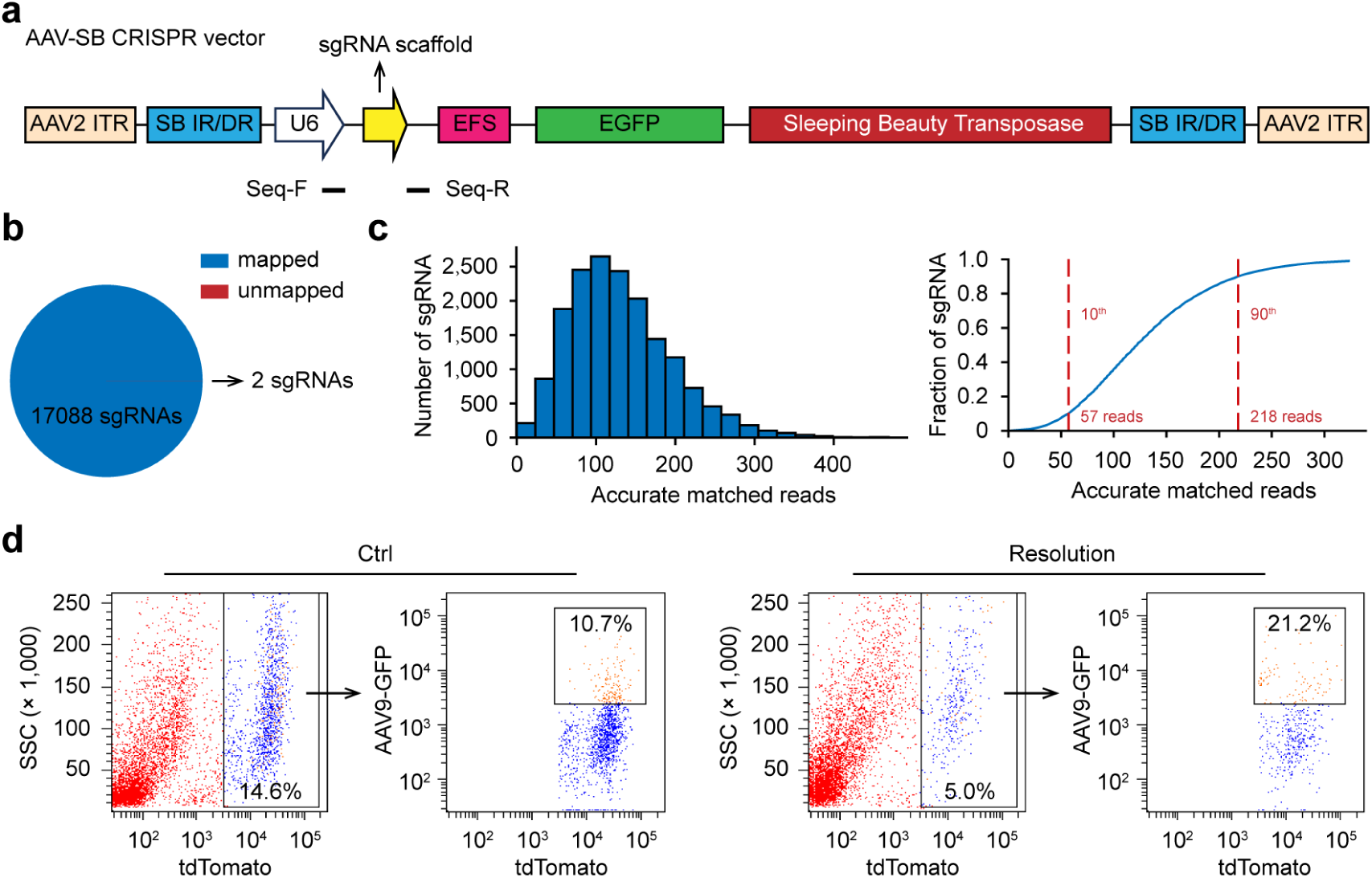
Generation of AAV9-SB-ME CRISPR library. (**a**) Schematic diagram of AAV-SB CRISPR vector containing the Sleeping Beauty transposon system and sgRNA scaffold. (**b**) sgRNA coverage in the AAV9-Metabolic CRISPR library. (**c**) The distribution of sgRNA sequencing read counts in the AAV9-Metabolic CRISPR library. (**d**) Flow cytometry analysis of GFP intensity in AECIIs from *Sftpc*-Cas9 mice transfected with the AAV9-Metabolic CRISPR library (5×10^11^ viral genome per mouse) through the trachea for two weeks followed by IAV infection (50 PFU). Ctrl, no infection; Resolution, 12 days post-IAV infection. Data are representative of three independent experiments (**d**).

To deliver this library into mice for screening, we packaged the vector library into AAV9 serotype virus, ultimately producing an AAV9-Metabolic viral library with a high titer of 2.42 × 10^13^ viral genome copies per milliliter (2.42 × 10^13^ vg/ml). To specifically identify essential metabolic genes for inflammation resolution in lung epithelial cells, we hybridized Rosa26-LSL-Cas9-tdTomato mice with *Sftpc*-Cre mice to generate animals expressing Cas9 and tdTomato in type II alveolar epithelial cells (*Sftpc*-Cas9), and injected the AAV9-Metabolic viral library into these *Sftpc*-Cas9 mice through the trachea (5 × 10^11^ vg/mouse). After two weeks, we detected the infection efficiency of the AAV9-Metabolic viral library both before IAV infection and at day 12 post-IAV infection, which is a resolution phage (indicated by body weight recovery and AECII proliferation) as previously described^24^. We found that nearly 10% of AECIIs (tdTomato-positive cells) were infected with the AAV9-Metabolic viral library (GFP-positive) in non-infected control mice, while more than 20% of AECIIs were infected during resolution of inflammation after IAV infection (**Fig. 1d**), suggesting that this library was suitable for functional screening.

### *In vivo* CRISPR screen of metabolic genes for resolving lung inflammation

AECIIs are reported to proliferate to promote tissue repair during inflammation resolution after IAV infection^25, 26^ (**Fig. S1a**). Thus, we reasoned that genes essential for inflammation resolution in these cells would be enriched after IAV infection (in “resolving cells”) compared to cells in the resting stage. To identify essential metabolic genes for inflammation resolution in AECIIs, we intratracheally injected *Sftpc*-Cas9 mice with the AAV9-Metabolic CRISPR library. Two weeks post-library transduction, we infected these mice with IAV, and then collected AECIIs at the inflammation resolution stage (12 days post-IAV infection) for sgRNA sequencing (**Fig. 2a**). Although *Sftpc*-Cas9 mice were susceptible to IAV-induced death, we ultimately obtained 2.1 × 10^6^ resting AECII cells as the control sample (more than 120 × coverage), and 9 × 10^5^ resolving AECII cells as the screen sample (more than 50 × coverage) for sgRNA sequencing. The results showed a sufficient mapping ratio and even distribution of sgRNA reads in control and screen samples (**Fig. S1b**, **c**), and almost all the nontargeting sgRNAs followed a linear regression line between control and screen samples (**Fig. S1d**), suggesting that this screening data was reliable. We performed MAGeCK analysis of sgRNA-targeted genes and found that sgRNAs targeting 151 genes were depleted, while sgRNAs targeting 147 genes were enriched in resolving AECII cells, suggesting these genes were either positively or negatively correlated with inflammation resolution in the lung (**Fig. 2b, c**).

**Figure 2.**
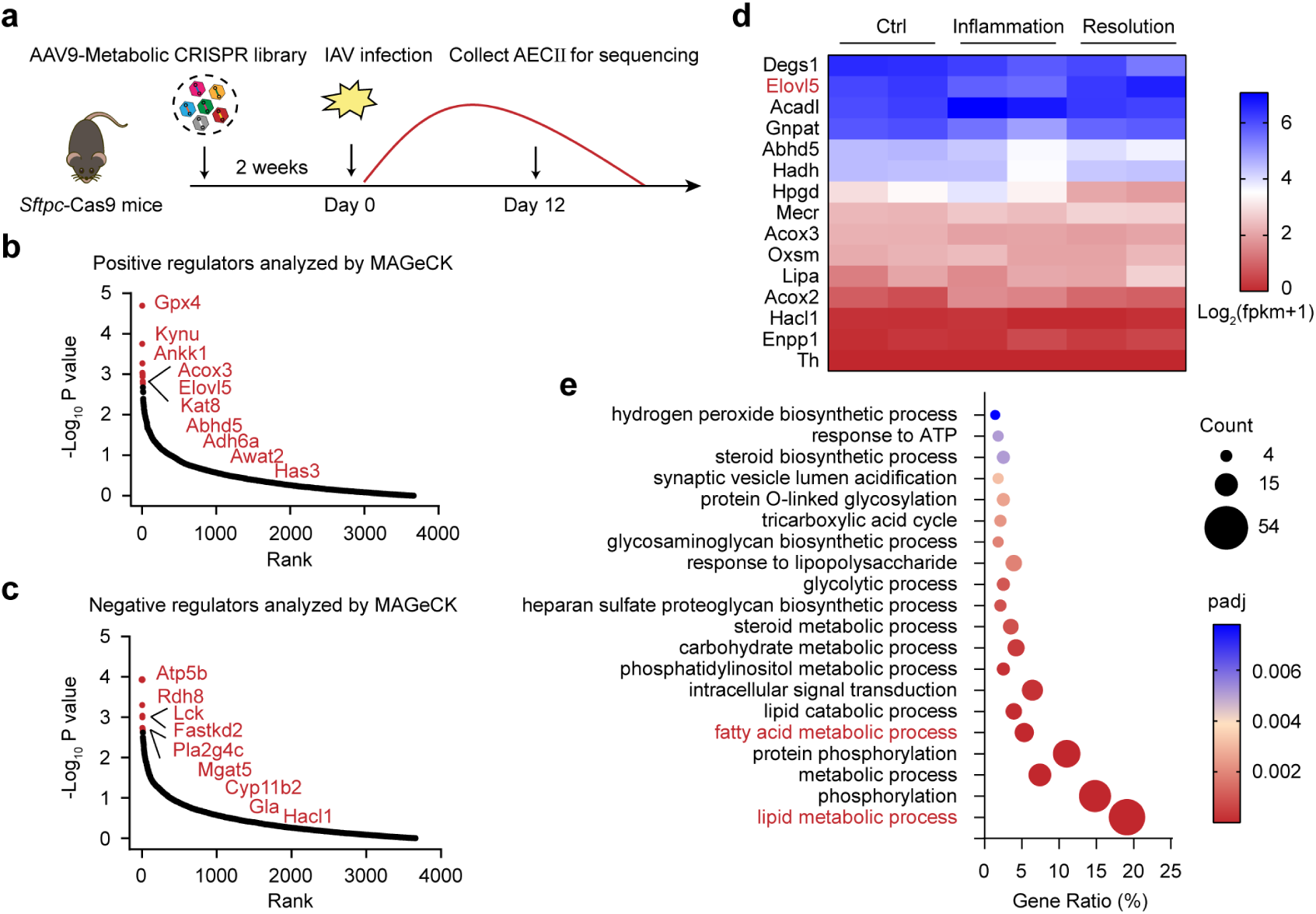
Identification of essential metabolic genes for lung inflammation resolution by *in vivo* CRISPR screening. (**a**) Schematic overview of *in vivo* CRISPR screen using AAV9-Metabolic CRISPR library in *Sftpc*-Cas9 mice. (**b**, **c**) Plots showing positive regulators (**b**) or negative regulators (**c**) for lung inflammation resolution identified in the screen by MAGeCK score analysis of sgRNA-targeted genes. Top 10-ranked genes are indicated in red color. (**d**) GO term analysis of differentially depleted and enriched genes in MAGeCK score analysis. (**e**) Heatmap showing expression of differentially depleted and enriched genes in fatty acid metabolism in AECIIs from mice at different times post-IAV infection (50 PFU). Ctrl, no infection; Inflammation, 7 days post-infection; Resolution, 12 days post-infection.

GO analysis of these depleted or enriched genes showed that lipid metabolic processes were highly enriched (**Fig. 2d**). This is reasonable since lipids, especially fatty acids, have been found to participate in the regulation of inflammation and immune response^27, 28^. Of note, enzymes involved in PUFA metabolism, including the elongase *Elovl5* and oxidases *Gpx4* and *Acox3*, were highly ranked among positive regulators (**Fig. 2b**), further supporting the idea that PUFA metabolism in AECIIs is essential for lung inflammation resolution. Another highly enriched GO term was protein phosphorylation (**Fig. 2d**). Unlike the lipid metabolism genes mentioned above, lymphocyte-specific protein tyrosine kinase (*Lck*), FAST kinase domain-containing protein 2 (*Fastkd2*), and cytosolic phospholipase A2 (*Pla2g4c*) were top-ranked in the list of genes that are downregulated during inflammation resolution (**Fig. 2c**), indicating phosphorylation in AECIIs may largely participate in promoting lung inflammation.

We focused on the fatty acid metabolic pathway and examined the expressions of these fatty acid metabolic genes in AECIIs during different stages of inflammation after IAV infection. We found that the expression of *Elovl5* was decreased during inflammation (7 days post-IAV infection) and recovered during the resolution stage (12 days post-IAV infection) (**Fig. 2e**), indicating that *Elovl5* might be involved in inflammation resolution and tissue repair. Indeed, *Elovl5* was one of the top 5 positive regulators, with two sgRNAs targeting different regions being identified by our screen (FDR < 0.05) (**Fig. 2b and Fig. S1e**). That two sgRNAs give the same outcome indicates that *Elovl5,* rather than an off-target, mediates the effect we observed. Therefore, *Elovl5* may be an essential metabolic gene in AECIIs for lung inflammation resolution.

### ELOVL5 promotes AECII-mediated lung inflammation resolution

To validate the screening results, we used an AECII-specific serotype of AAV, AAV- LungM3, to deliver an *Elovl5*-targeting sgRNA into CAG-Cas9 mice, thus knocking down *Elovl5* expression in AECIIs. We found that AAV-LungM3 specifically infected AECII cells *in vivo*, and *Elovl5* sgRNA significantly decreased the expression of *Elovl5* in the lungs of CAG-Cas9 mice (**Fig. S2a, b**). We then infected these mice with IAV and found that compared to the non-targeting control sgRNA, *Elovl5* sgRNA-transfected mice were more susceptible to IAV-induced death (**Fig. 3a**). We further examined lung inflammation of these mice in the resolution stage (10 days post-IAV infection). We found that the total protein level in bronchoalveolar lavage fluid (BALF) was higher in *Elovl5* sgRNA-transfected CAG-Cas9 mice than in control mice (**Fig. 3b**), indicating that *Elovl5* sgRNA-transfection inhibited lung epithelium repair in CAG-Cas9 mice. In addition, the expression level of inflammation-resolving cytokines, including *Il10* and *Tgfb2*, was decreased in *Elovl5* sgRNA-transfected mice. Meanwhile, expression levels of the proliferation marker *Ki-67* and AECII marker *Sftpc* were also significantly decreased (**Fig. 3c**). Consistently, *Elovl5* sgRNA-transfected CAG-Cas9 mouse lung tissue showed less Ki-67 and SP-C staining, and more severe lung inflammation (**Fig. 3d, e**). These data suggested that *Elovl5* knockdown in AECII inhibited inflammation resolution and tissue repair in CAG-Cas9 mice after IAV infection.

**Figure 3.**
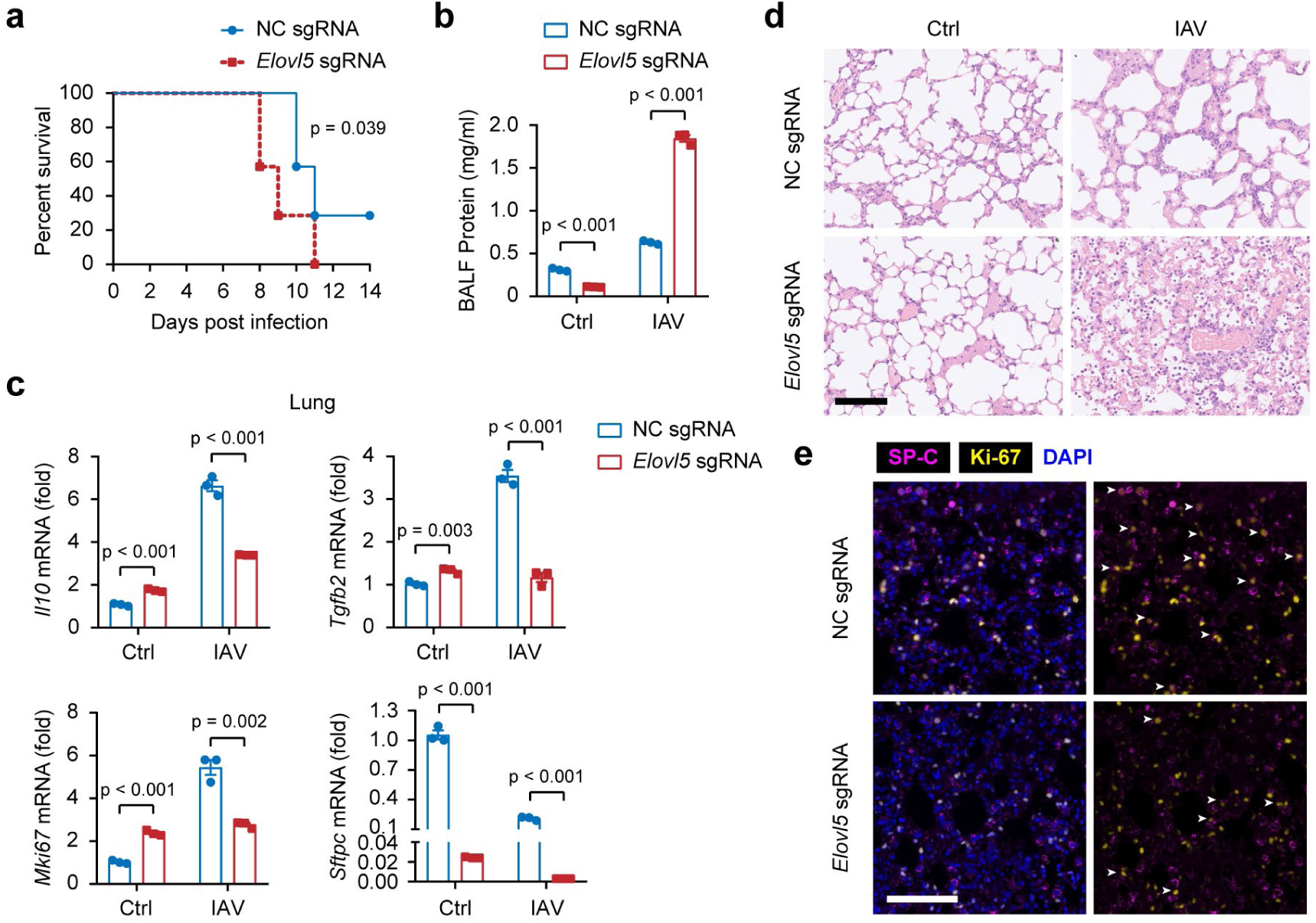
ELOVL5 promotes lung inflammation resolution in CAG-Cas9 mice after IAV infection. (**a**) Survival curves of CAG-Cas9 mice at different times post-IAV infection (50 PFU) after transfecting with AAV-LungM3 containing negative control sgRNA (n = 8) or sgRNA targeting *Elovl5* (n = 7) (5 × 10^10^ viral genome copies per mouse) for two weeks. (**b**) BCA analysis of protein level in BALF from CAG-Cas9 mice at day 10 post-IAV infection (50 PFU) after transfecting with AAV-LungM3 containing negative control sgRNA or sgRNA targeting *Elovl5* (5 × 10^10^ viral genome copies per mouse) for two weeks. (**c**) qPCR analysis of *Il10*, *Tgfb2*, *Mki67* and *Sftpc* mRNA expressions in lungs from mice described in **b**. (**d**) Hematoxylin and eosin staining of lung sections from the mice described in **b**. Scale bar, 100 μm. (**e**) Immunofluorescence staining of Ki-67 and SP-C in lung sections from mice described in **b**. Scale bar, 100 μm. White arrows represent Ki-67^+^ AECIIs. Data were pooled from two independent experiments (**a**) or representative of three independent experiments (**b**-**e**; shown as mean ± SD in **b**, **c**), Log-rank (Mantel-Cox) analysis in **a**, unpaired Student’s t test in **b**, **c**.

To further investigate the function of ELOVL5 in AECIIs and to exclude the effects caused by the Cas9 protein, we used AAV-LungM3-shRNA to deplete the expression of *Elovl5* in AECIIs in wild-type C57BL/6 mice (**Fig. S2c**). We found that *Elovl5* depletion dampened the body weight recovery of mice after IAV infection (**Fig. 4a**). The protein level and the numbers of neutrophils and macrophages in BALF were higher in *Elovl5*-depleted mice compared to control mice (**Fig. 4b, c, Fig. S2d, e**) at day 12 post-IAV infection, suggesting that *Elovl5* depletion in AECIIs inhibited the resolution of pulmonary inflammation and repair of epithelial leakage. Furthermore, *Elovl5*-depleted mice showed increased lung inflammation and decreased Ki-67 staining in AECIIs (**Fig. 4d-f, Fig. S2f**), suggesting that *Elovl5* depletion disrupted inflammation resolving responses in AECII cells. Together, these results demonstrate that the metabolic gene *Elovl5* in AECIIs is essential for lung inflammation resolution after IAV infection.

**Figure 4.**
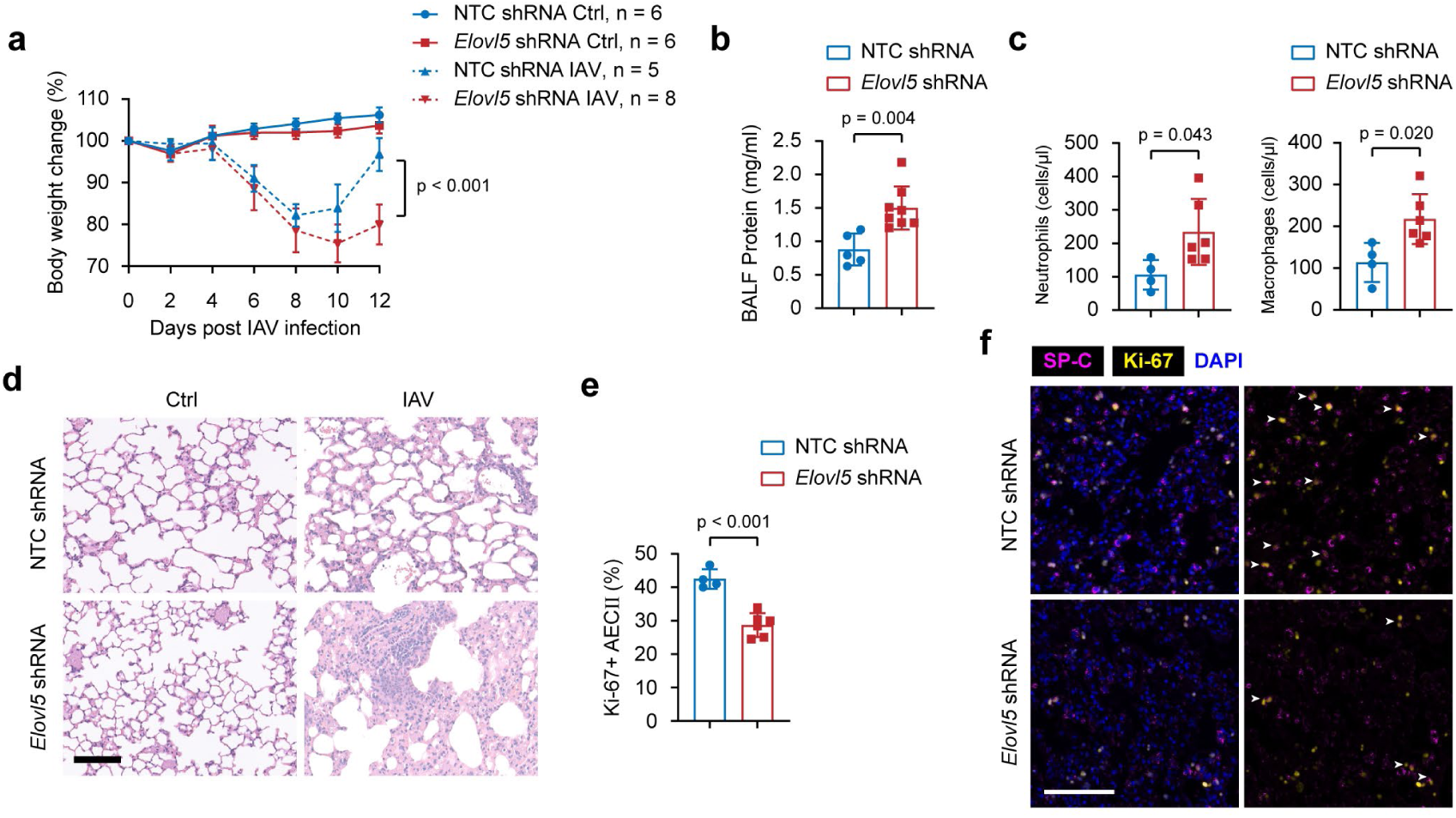
ELOVL5 promotes lung inflammation resolution in C57BL/6 mice after IAV infection. (**a**) Body weight change of C57BL/6 mice at different times post-IAV infection (50 PFU) after transfecting with AAV-LungM3 containing non-targeting control shRNA or shRNA targeting *Elovl5* (5 × 10^10^ viral genome copies per mouse) for two weeks. (**b**) BCA analysis of protein level in BALF from C57BL/6 mice at day 12 post-IAV infection (50 PFU) after transfecting with AAV-LungM3 containing non-targeting control shRNA or shRNA targeting *Elovl5* (5 × 10^10^ viral genome copies per mouse) for two weeks. (**c**) Flow cytometry analysis of neutrophil and macrophage numbers in BALF from mice described in **b**. (**d**) Hematoxylin and eosin staining of lung sections from the mice described in **b**. Scale bar, 100 μm. (**e**) Flow cytometry analysis of AECII proliferation from mice described in **b**. (**f**) Immunofluorescence staining of Ki-67 and SP-C in lung sections from mice described in **b**. Scale bar, 100 μm. White arrows represent Ki-67^+^ AECIIs. Data were pooled from two independent experiments (**a**, **b**) or representative of three independent experiments (**c**-**f**; shown as mean ± SD in **b**, **c**, **e**), Two-way ANOVA analysis in **a**, two-tailed unpaired Student’s t test in **b**, **c**, **e**.

### ELOVL5 interacts with STING to promote inflammation resolution

We went further to investigate how ELOVL5 promotes inflammation resolution in AECIIs. We used AAV-LungM3-shRNA to deplete the expression of *Elovl5* in AECIIs *in vivo*, and then isolated these AECIIs at day 12 post-IAV infection. We found that expression of the inflammatory cytokine *Il1b* was increased in *Elovl5*-depleted AECIIs, while expression of the resolving cytokine *Il10* and genes involved in the proliferation of AECIIs after injury, including *Ctgf* and *Cyr61*^13, 14^, were decreased (**Fig. 5a**). Next, we generated *Elovl5*-deficient mouse lung epithelial cells (MLE-12) *in vitro* using the CRISPR-Cas9 system (**Fig. S3a**), and found that *Elovl5* deficiency increased expression of the inflammatory cytokines *Il1b* and *Tnf* while decreasing expression of the proliferation genes *Ctgf* and *Cyr61* in MLE-12 cells after IAV infection for 24 h (**Fig. 5b**). These data suggest that ELOVL5 inhibited the inflammatory response and promoted proliferation of AECIIs after viral infection.

**Figure 5.**
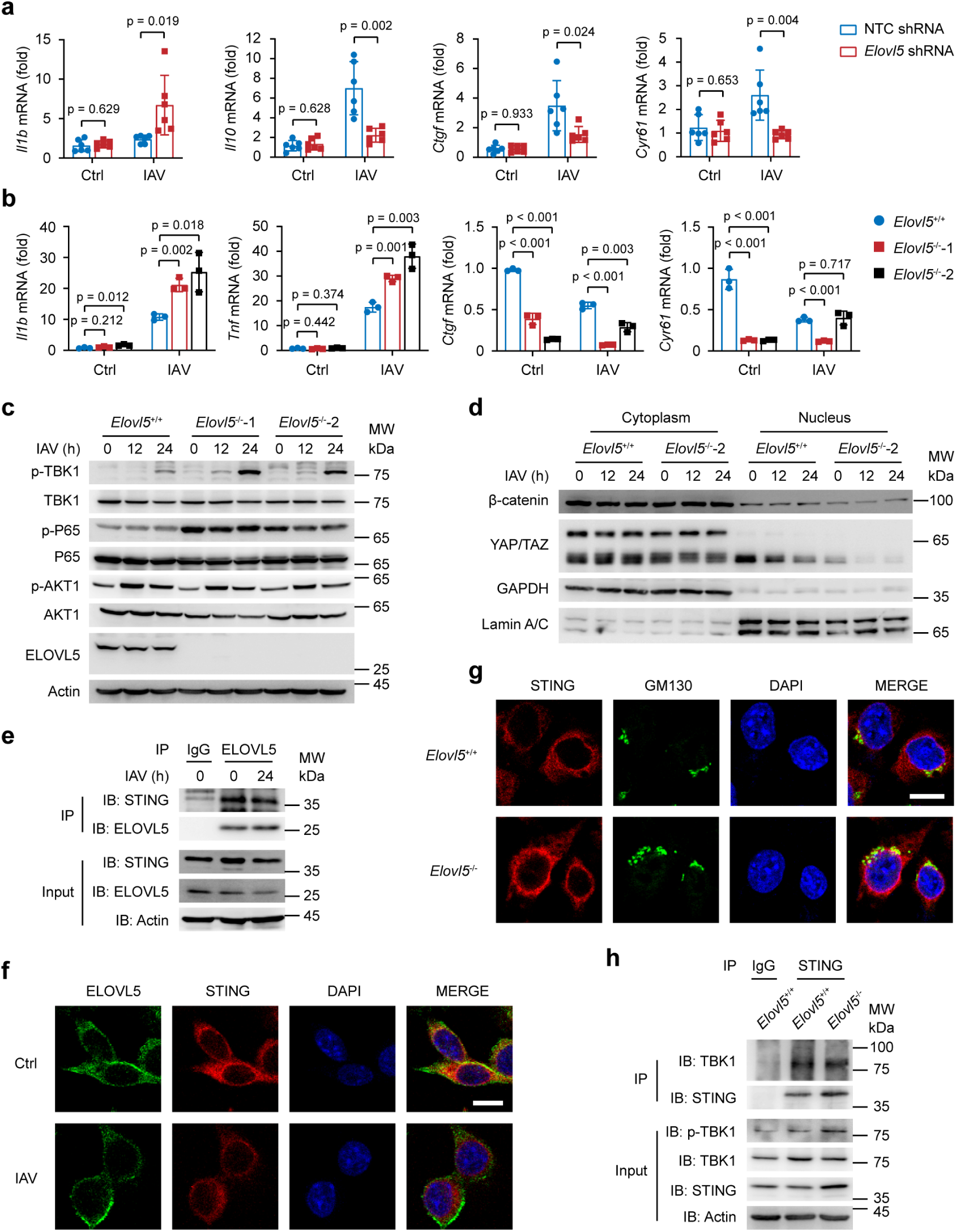
ELOVL5 promotes inflammation resolution through interacting with STING in mouse lung epithelial cells. (**a**) qPCR analysis of *Il1b*, *Il10*, *Ctgf* and *Cyr61* mRNA expressions in AECIIs from mice at day 12 post-IAV infection (50 PFU) after transfecting with AAV-LungM3 containing non-targeting control shRNA or shRNA targeting *Elovl5* (5 × 10^10^ viral genome copies per mouse) for two weeks. (**b**) qPCR analysis of *Il1b*, *Tnf*, *Ctgf* and *Cyr61* mRNA expression in *Elovl5*^+/+^ and *Elovl5*^-/-^ MLE-12 cells infected with IAV (MOI = 1) for 24 h. (**c**) Immunoblot analysis of signaling pathway components in *Elovl5*^+/+^ and *Elovl5*^-/-^ MLE-12 cells infected with IAV (MOI = 1) for indicated times. (d) Immunoblot analysis of nuclear translocation of β-catenin and YAP/TAZ in *Elovl5*^+/+^ and *Elovl5*^-/-^ MLE-12 cells infected with IAV (MOI = 1) for indicated times. (e) Immunoblot analysis of interaction between ELOVL5 and STING in MLE-12 cells infected with IAV (MOI = 1) for 24 h. (**f**) Immunofluorescence analysis of interaction between ELOVL5 and STING in MLE-12 cells infected with IAV (MOI = 1) for 24 h. Scale bar, 10 μm. (**g**) Immunofluorescence analysis of interaction between STING and GM130 in *Elovl5*^+/+^ and *Elovl5*^-/-^ MLE-12 cells infected with IAV (MOI = 1) for 24 h. Scale bar, 10 μm. (**h**) Immunoblot analysis of interaction between STING and TBK1 in *Elovl5*^+/+^ and *Elovl5*^-/-^ MLE-12 cells infected with IAV (MOI = 1) for 24 h. Data were pooled from two independent experiments (**a**) or representative of three independent experiments (**b**-**h**; shown as mean ± SD in **a**, **b**), unpaired Student’s t test in **a**, **b**.

To demonstrate how ELOVL5 regulates the expression of these genes, we examined the signaling pathways involved in inflammation and AECII proliferation. We found that the phosphorylation of TBK1 was increased in *Elovl5*-deficient MLE-12 cells, while the phosphorylation of AKT1 and the nuclear translocation of β-catenin and YAP/TAZ, which are critical for AECII proliferation after injury, were decreased (**Fig. 5c, d, Fig. S3b**).

It has been found that the influenza virus activates STING, which is an essential adaptor for activating TBK1^29, 30^. STING undergoes translocation from the ER to Golgi^31, 32, 33^, then recruits and activates TBK1 after viral infection^34^. ELOVL5 is a transmembrane protein located in the endoplasmic reticulum (ER), therefore, we wondered whether ELOVL5 might interact with STING to promote inflammation resolution and tissue repair. Using immunoprecipitation and immunofluorescence assays, we confirmed that ELOVL5 interacted with STING in MLE-12 cells, and their interaction was decreased after IAV infection (**Fig. 5e, f, Fig. S3c, d**). We next assessed whether ELOVL5 affected the translocation of STING and the interaction between STING and TBK1. We found that *Elovl5* deficiency increased the co-localization of STING and the Golgi marker GM130, and the interaction between STING and TBK1 was also increased in *Elovl5*-deficient MLE-12 cells after IAV infection (**Fig. 5g, h**). These data suggest that ELOVL5 promotes inflammation resolution in AECIIs through binding with STING and inhibiting STING-TBK1 interaction.

### ELOVL5 decreases eicosanoid levels in AECII for lung inflammation resolution

As ELOVL5 is a PUFA elongase, and PUFAs are important sources for generating SPMs, we next investigated whether PUFAs and PUFA-related metabolites participate in ELOVL5-mediated inflammation-resolving response in AECIIs. We pretreated *Elovl5*-deficient MLE-12 cells with arachidonic acid (AA) or eicosapentaenoic acid (EPA), which are downstream products of ELOVL5, before IAV infection (**Fig. 6a**). We found that treatment with AA or EPA reversed the IAV infection-increased expression of inflammatory cytokines caused by *Elovl5* deficiency (**Fig. 6b**), suggesting that the inflammation-resolving effect of *Elovl5* depends on these PUFAs. However, pretreatment with linoleic acid (LA) or α-linolenic acid (ALA), which are substrates of ELOVL5 that generate PUFAs, also reversed the pro-inflammatory effect of *Elovl5* deficiency (**Fig. 6c**). These results prompted us to investigate whether another LA or ALA-linked metabolic pathway participated in ELOVL5-mediated inflammation resolution in AECIIs. Then, we analyzed eicosanoids to examine the levels of PUFA-related lipid mediators in *Elovl5*-deficient MLE-12 cells in the absence or presence of ALA after IAV infection. We found that *Elovl5* deficiency significantly increased the prostaglandins (PGs), the lipid mediators of inflammation, including PGE2, PGD2, PGF2a and PGF2b in MLE-12 cells after IAV infection (**Fig. 6d**). However, pretreatment with ALA significantly decreased the level of these inflammatory lipid mediators (**Fig. 6e**), suggesting that ALA might reverse *Elovl5* deficiency-induced inflammation via decreasing these mediators. In addition to PGs, levels of other intermediators, including hydroxytetraenoic acid (HETE), hydroxyl eicosapentaenoic acid (HEPE) and hydroxy docosahexaenoic acid (HDHA), which are produced during eicosanoid class-switching^35^, were also increased in *Elovl5*-deficient MLE-12 cells and decreased after ALA treatment (**Fig. 6d, e**). These data suggest that ELOVL5 decreases eicosanoids in AECIIs to prevent inflammatory cytokine expression, thereby promoting inflammation resolution and tissue recovery.

**Figure 6.**
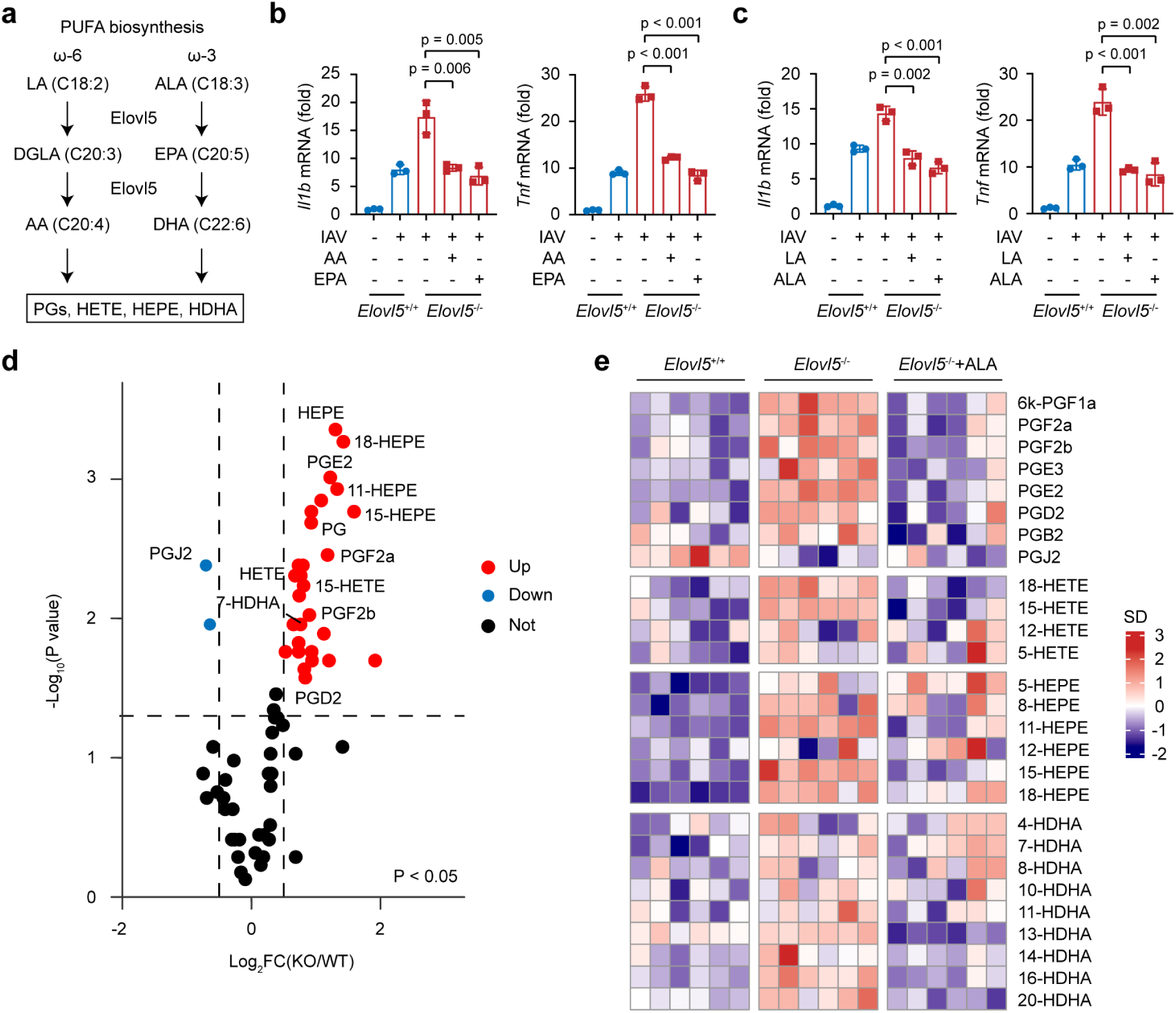
ELOVL5 decreases eicosanoid levels to promote inflammation resolution in mouse lung epithelial cells. (**a**) Schematic diagram showing PUFA and eicosanoid biogenesis. (**b**) qPCR analysis of *Il1b* and *Tnf* mRNA expressions in *Elovl5*^+/+^ and *Elovl5*^-/-^ MLE-12 cells pretreated with AA (20 μM) or EPA (20 μM) for 12 h, followed by IAV infection (MOI = 1) for 18 h. (**c**) qPCR analysis of *Il1b* and *Tnf* mRNA expressions in *Elovl5*^+/+^ and *Elovl5*^-/-^ MLE-12 cells pretreated with LA (20 μM) or ALA (20 μM) for 12 h, followed by IAV infection (MOI = 1) for 18 h. (**d**) Volcano plots showing differentially enriched eicosanoids in *Elovl5*^+/+^ and *Elovl5*^-/-^ MLE-12 cells infected with IAV (MOI = 1) for 24 h. |log2Fold Change| > 0.5, p < 0.05. (**e**) Heatmap showing differential eicosanoid levels in *Elovl5*^+/+^, *Elovl5*^-/-^ or *Elovl5*^-/-^ MLE-12 cells pretreated with ALA (20 μM) for 12 h, followed by IAV infection (MOI = 1) for 24 h. SD value represents the standard deviation of each metabolite to the average. Data are representative of three independent experiments (**b**, **c**; shown as mean ± SD in **b**, **c**), unpaired Student’s t test in **b**, **c**.

## Discussion

In this study, we systematically searched for metabolic genes that regulate lung inflammation resolution. Through an *in vivo* CRISPR screen targeting 2682 mouse metabolic genes, we identified ELOVL5 as an essential metabolic enzyme that mediates a pro-resolving response as a dual immune-metabolic regulator in alveolar epithelial cells after influenza virus infection. ELOVL5 expression is dynamically decreased at the early stage of IAV infection, whereas up-regulated in AECIIs at the resolving stage, which promotes inflammation resolution and AECII proliferation for tissue repair. On the one hand, ELOVL5 could maintain cellular PUFAs homeostasis and decrease inflammatory lipid mediator accumulation as a metabolic enzyme. More importantly, we found a previously uncharacterized role of ELOVL5 in sequestering STING and suppressing STING-mediated inflammatory signaling in the late stage of infection. Our results broaden the potential to manipulate ELOVL5, PUFA biogenesis, and eicosanoid levels for the treatment of lung inflammation-related diseases.

Studies on immunometabolism have provided evidence for lipid metabolism-mediated immune response regulation, however, the specific enzymes involved, and the complex interplay between immune signaling and metabolism pathways in distinct cell types require further investigation. Here we uncovered that a fatty acid elongase, ELOVL5, could mediate lung inflammation resolution in AECIIs. Although ELOVL5 is an elongase that generates long-chain polyunsaturated fatty acids, we found that *Elovl5* deficiency increased eicosanoid levels, indicating the complicated regulatory metabolic network during inflammation resolution. Interestingly, supplement with ELOVL5 upstream and downstream fatty acids both reversed increased inflammatory cytokine expressions caused by *Elovl5* deficiency. It will be worthy to further investigate how *Elovl5* deficiency increased eicosanoid levels, and how ALA pretreatment decreased eicosanoid levels. One possibility is that ELOVL5 might function in promoting lipid mediator class switch^35^ during inflammation resolution. On the other hand, eicosanoids are derived from the oxidation of PUFAs by lipoxygenase, cyclooxygenase and cytochrome P450 during inflammation^35^, *Elovl5* deficiency might partially contribute to the excessive oxidation of PUFAs through excess ROS generation^36^, which needs to be further investigated.

As a central hub of innate immune response and inflammation, the activation and suppression of STING must be tightly regulated. A STING ER exit protein (STEEP) has been reported to interact with STING to promote ER-to-Golgi trafficking^32^. Besides, tyrosine kinase HER2 could associate with STING and recruit AKT1 to prevent STING-TBK1 interaction^37, 38^. However, whether STING could be regulated by metabolic enzymes and metabolites remains elusive. Many metabolic enzymes have been discovered by their moonlighting function other than catalytic activity in metabolism. Here we found beyond keeping fatty acid homeostasis, ELOVL5 could also act as a checkpoint of STING pathway by directly interacting with STING and sequestering STING from the Golgi during viral infection. Further studies are needed to fully clarify the relationship between ELOVL5 and ALA-eicosanoids metabolic axis, as well as the relationship between eicosanoids and STING signaling. Our results demonstrate a previously undescribed mechanism by which the metabolic enzyme, lipid metabolites and immune signaling coordinate to regulate active inflammation resolution and tissue repair in a specific cell type, providing more evidence for understanding immune-metabolic crosstalk and developing drug targets in controlling inflammatory diseases.

Unresolved lung inflammation leads to various diseases, such as sepsis, fibrosis, autoimmune diseases in addition to virus infection^39, 40^. Here we found that *Elovl5* knockdown in AECIIs significantly increased inflammatory cell infiltration and epithelial leakage, decreased AECII proliferation in the lung of mice at the resolution stage of IAV infection, suggesting a protective role of ELOVL5 in inflammatory diseases. Besides, PUFA supplement has been widely studied in many diseases^41^, therefore, it will be intriguing to investigate whether manipulating ELOVL5, in combination with PUFA supplement would broadly benefit the treatment of other inflammatory disorders.

AECIIs play important regulatory roles in pulmonary immune response and tissue repair. Our results provide evidence that AECIIs could upregulate a fatty acid elongase to reprogram lipid metabolism and innate signaling to ultimately terminate inflammatory response and initiate cell proliferation in the resolution stage of IAV infection. Single-cell RNA sequencing data have identified that AECII subsets with high Wnt signaling and *Ctgf* expression have more potential to proliferate and differentiate^13, 42^. We found that ELOVL5 promoted the nuclear translocation of β- catenin and the expression of *Ctgf*, suggesting that ELOVL5 could participate in maintaining the “stemness” of AECIIs, and thus promote lung regeneration after injury. SPMs promote the proliferation and differentiation of AECIIs^43^, our results further linked fatty acid metabolism to AECII function. Future studies remain to be done to demonstrate whether and how ELOVL5-mediated lipid metabolism, including PUFAs and eicosanoids, affects AECII “stemness”.

CRISPR screen has been widely used to identify functional targets in different disease models, including autoimmune diseases, infections and tumors^44, 45, 46^. The major difficulty to discover essential regulators for inflammation resolution is to manipulate target genes within the tissue microenvironment. Here by applying AAV9- SB-ME CRISPR library into AECIIs-specifically expressing Cas9 mice, we successfully screened essential metabolic genes for AECII-mediated lung inflammation resolution, suggesting that this system was ideal for identifying metabolic regulators *in vivo*. Therefore, future studies could focus on these metabolic regulators, and by using different serotypes of AAV, this system will also be promising in identifying essential metabolic mediators in different disease models via targeting different tissue cells.

Except for ELOVL5, other very long-chain fatty acid elongase family members have also been implicated in regulating immune response. For example, *Elovl6* has been found to hamper central nervous system repair through inhibiting a repair-promoting phenotype in foamy macrophages^47^. *Elovl1* has been found to be a target to sustain effector functions and memory phenotypes of CD8^+^ T cells^48^, suggesting the broader regulating function of this family in different cells within their respective tissue microenvironment. Of note, *Elovl5* has been found to be up-regulated to promote ferroptosis sensitization in mesenchymal cancer cells^49, 50^. Ferroptosis is closely related to inflammation and whether ferroptosis is involved in *Elovl5*-mediated regulation of inflammation resolution needs to be investigated in the future. Fully demonstrating the function and mechanism of ELOVL5 and other family enzymes will provide further insight for developing effective PUFA-based therapies under different immunopathological conditions.

## Materials and Methods

### Mice

Rosa26-LSL-Cas9-tdTomato mice, *Sftpc*-Cre mice, CAG-Cas9 mice were obtained from GemPharmatech Co. Ltd, Nanjing, China. C57BL/6 mice were from Beijing Vital River Laboratory, Beijing, China. All mice were maintained in specific pathogen-free conditions. For AAV transduction, both female and male, 8-week-old mice were used. Mice were randomly assigned to different treatment groups. All mouse experiments were performed under the supervision of the Institutional Animal Care and Use Committee, Institute of Basic Medical Sciences, Chinese Academy of Medical Sciences, Beijing, China (ACUC-A01-2022-013).

### Cell lines

Mouse lung epithelial cells (MLE-12 cells) and human embryonic kidney cells (HEK293T cells) were from ATCC. *Elovl5*-deficient MLE-12 cells were generated using a guide RNA targeting exon 6 of mouse *Elovl5*: 5′-CGTCCATGCGTCCCTACCTC -3′.

### Plasmids, reagents and pathogens

AAV-CRISPR vector was generated by cloning Sleeping Beauty transposase and sgRNA scaffold into the AAV vector. Flag-ELOLV5 plasmid was generated by cloning full-length mouse ELOLV5 into pcDNA3.1 plasmid. Arachidonic acid (HY-109590), eicosapentaenoic acid (HY-B0660), linoleic acid (HY-N0729) and α-linolenic acid (HY-N0728) were from MedChemExpress. Influenza virus strain A/Puerto Rico/8/1981 H1N1 (PR8) was amplified as previously described^24^.

### Design and generation of AAV9-Metabolic CRISPR library

The search for metabolic genes was based on metabolism pathway genes in the KEGG database (https://www.genome.jp/kegg/pathway.html) and Recon 2 database (http://humanmetabolism.org), 2682 mouse metabolic genes were selected. 6 sgRNAs for each metabolic gene (4 sgRNAs for *Uxs1*) and 1000 nontargeting sgRNAs were designed to generate a metabolic sgRNA library which contained 17090 sgRNAs in total (**Data S1**). The metabolic sgRNA library was then synthesized and cloned into the AAV-SB vector to generate an AAV-Metabolic CRISPR vector library by Gibson Assembly. The coverage and representation of sgRNAs in the vector library were verified by next-generation sequencing.

The vector library was packaged into AAV9 serotype viruses to produce an AAV9-Metabolic CRISPR library. The virus titer was measured by quantitation of the number of viral genomes using the ITR sequence.

### AAV9-Metabolic CRISPR screen in a mouse model of influenza virus infection

To screen essential metabolic genes for lung inflammation resolution, 5 × 10^11^ viral genome AAV9-Metabolic CRISPR library was transduced into the lungs of *Sftpc*-Cas9 mice via the trachea. After two weeks, mice were intranasally infected with influenza virus (50 PFU in PBS). The body weight of mice was then monitored every day. 12 days post-viral infection, 9 × 10^5^ AECIIs from mice at the inflammation resolution stage were sorted as the experimental sample (> 50 × coverage), while 2.1 × 10^6^ resting AECII cells from non-infected mice were sorted as control samples (> 120 × coverage). Genomic DNA was extracted from samples using the DNeasy Blood & Tissue Kit (Qiagen). The sgRNA sequence in genomic DNA was amplified by PCR using primers: F: 5’-AATGGACTATCATATGCTTACCGTAACTTGAAAGTA -3’, R: 5’-CAATTCCCACTCCTTTCAAGACCTAGTCGACGTTTA -3’. PCR products were purified using the QIAquick PCR Purification Kit (Qiagen). DNA was recovered for concentration measurement and next-generation sequencing on the Illumina platform. Sequencing results were deposited in NCBI GEO database under accession code GSE298405.

### Type II alveolar epithelial cell isolation

Mouse primary type II alveolar epithelial cells (AECIIs) were isolated as previously described^24^. Briefly, mouse lung was intratracheally injected with 1 mL Dispase II (50 U/mL) (Roche) and digested for 45 min at room temperature. Lung cells were harvested, and CD45^-^EPCAM^+^MHCII^+^ cells were labeled and sorted as AECIIs.

### AAV-LungM3 transduction and influenza virus infection mouse model

AAV-LungM3 carrying the *Elovl5* sgRNA (CGTCCATGCGTCCCTACCTC) or the non-targeting control sgRNA (GCACTACCAGAGCTAACTCA), *Elovl5* shRNA (GCCCAGAGCTTGTTAGTTTAA) or non-targeting control shRNA (CCTAAGGTTAAGTCGCCCTCG) were generated by Obio Technology (Shanghai, China). 5 × 10^10^ viral genome copies diluted in 50 μL sterile saline were injected into each mouse lung via trachea. After two weeks, mice were intranasally infected with influenza virus (50 PFU in PBS). Body weight and survival were monitored every day. 12 days post-viral infection, BALF was collected for protein concentration and cell infiltration analysis; lung tissue was collected for AECII isolation and histopathological examination, including hematoxylin-and-eosin staining and immunofluorescence staining of Ki-67 and SP-C using anti-Ki-67 (D3B5) from Cell Signaling Technology and anti-SP-C (EPR28174-18) from Abcam.

### Flow cytometry analysis

BALF cells or AECIIs were harvested and labeled with antibodies for 30 min at 4 °C. For intracellular Ki-67 staining, cells were fixed and permeabilized using FOXP3 Fix/Perm Buffer Set (BioLegend). After that, cells were washed with PBS and then analyzed by LSRFortessa or sorted by FACSAriaII (BD Biosciences). The following antibodies were used: BV421 anti-mouse CD45 (30-F11), APC anti-mouse Ep-CAM (G8.8), PE/Cy7 anti-mouse I-A/I-E (M5/114.15.2), PE anti-mouse Ki-67 (16A8), APC anti-mouse CD11b (M1/70), FITC anti-mouse Gr-1 (RB6-8C5), PE anti-mouse F4/80 (BM8) and Precision Count Beads, all from BioLegend.

### Quantitative PCR

Total RNA was extracted using Trizol (Thermo Scientific) or Direct-zol RNA Microprep Kits (Zymo Research), cDNA was synthesized using ReverTra Ace qPCR RT Master Mix (TOYOBO). Quantitative PCR (q-PCR) was performed using SYBR Green Realtime PCR Master Mix (TOYOBO). *Rpl32* was used as an internal control. Primers used were listed in **Table S1**.

### Transcriptome analysis

AECIIs were isolated from C57BL/6 mice at day 0, 7 or 12 post-IAV infection (50 PFU) separately. Total RNA of AECIIs was extracted using Direct-zol RNA Microprep Kits (Zymo Research). mRNAs were purified by oligo(dT) beads and libraries were generated using the NEBNext Ultra RNA Library Prep Kit. RNA-seq was performed on the Illumina platform. Clean reads were aligned to the reference genome using HISAT2, and featureCounts was used to count the reads mapped to each gene. The RNA sequencing results were deposited in NCBI GEO database under accession code GSE298405.

### Immunoprecipitation and immunoblot

For immunoprecipitation, cells were lysed in IP lysis buffer (Thermo Scientific) supplemented with protease inhibitor cocktail (Merck). After centrifugation, supernatants were collected and incubated with ELOVL5 antibody at 4 °C overnight. The next day, protein A/G magnetic beads were added for another 2 h’s incubation. ELOVL5-interacting proteins were eluted by boiling in SDS loading buffer. Total cell proteins were obtained using RIPA lysis buffer supplemented with protease inhibitor cocktail (Merck). Cytoplasm and nuclear proteins were separated using Minute Cytoplasmic and Nuclear Extraction Kits (Invent). After concentration measurement, protein samples were boiled in SDS loading buffer before immunoblot analysis. For ELOVL5 detection, protein samples were incubated with SDS loading buffer at room temperature. The following antibodies were used: anti-ELOVL5 (E-10) from Santa Cruz Biotechnology. Anti-p-TBK1 (D52C2), anti-TBK1 (E8I3G), anti-p-P65 (93H1), anti-P65 (D14E12), anti-p-AKT1 (D7F10), anti-AKT1 (C73H10), anti-STING (D2P2F), anti-Flag (D6W5B), anti-Lamin A/C (4C11), anti-β-catenin (6B3), anti-YAP/TAZ (D24E4) and normal rabbit IgG antibodies from Cell Signaling Technology. Anti-GAPDH (3H12) and anti-Actin (6D1) antibodies from MBL.

### Immunofluorescence

Cells were seeded on glass coverslips. After IAV infection, cells were fixed with 4% paraformaldehyde, permeabilized with 0.2% Triton X-100 in phosphate-buffered saline, followed by blocking with 1% BSA. Anti-ELOVL5 (E-10) (Santa Cruz Biotechnology), anti-STING (Alexa Fluor 647 Conjugate) (Cell Signaling Technology), anti-GM130 (BD Biosciences) primary antibodies and fluorescent dye-conjugated secondary antibodies (Thermo Scientific) were used. Nuclei were stained with DAPI. Fluorescence signals were detected and analyzed using a Leica TCS SP8 STED 3X.

### Eicosanoid analysis

Eicosanoids in samples were quantitated at LipidALL Technologies (China). Cells were extracted in a buffer comprising methanol containing 0.1% (w/v) of butylated hydroxytoluene and butylated hydroxyanisole with formic acid and internal standard cocktail added. Samples were vortexed to promote thorough mixing. A fixed amount of ceramic beads pre-cleaned with methanol was then added and the cell samples were incubated at 1500 rpm for 12 h at 4 °C to ensure efficient extraction of eicosanoids. The samples were centrifuged at 4 °C for 10 min at 12000 rpm, and the supernatant was extracted. The extraction was repeated for a second round. The pooled supernatants were enriched for eicosanoids via solid phase extraction (SPE) using Oasis Prime HLB columns (30 mg, Waters, USA). The internal standard cocktail contained PGD2-d4, PGE2-d4, PGF2a-d4, 6-keto-PGF1a-d4, 13,14-dihydro-15-keto prostaglandin D2-d4, 13,14-dihydro-15-keto prostaglandin F2a-d4, HETE: 5(S)-hydroxy-eicosatetraenoic acid-d8, HETE: 12(S)-hydroxy-eicosatetraenoic acid-d8, HETE: 15(S)-hydroxy-eicosatetraenoic acid-d8, DiHOME: 9,10-dihydroxy octadecenoic acid-d4, DiHOME: 12,13-dihydroxy octadecenoic acid-d4, resolvin D1-d5, leukotriene B4-d4, HODE: 9(S)-hydroxy-octadecadienoic acid-d4, HODE: 13(S)-hydroxy-octadecadienoic acid-d4, ARA-d11, d31-16:0, EOME: 9,10-EOME-d4, DHET: 11,12-DHET-d10, in methanol (Cayman chemicals, USA). SPE eluents were transferred to tubes containing 20 µL of ethanol:glycerol 1:1 (v/v) to prevent complete desiccation, and dried under flowing stream of nitrogen gas. The dried extract was re-constituted immediately in 50 µL of water:acetonitrile:formic acid 63:37:0.02 (v/v/v) for mass spectrometric analysis. Eicosanoid analysis were conducted on an Shimadzu 40X3B-UPLC coupled to Sciex QTRAP 6500 Plus (Sciex, USA). Eicosanoids were separated on a Phenomenex Kinetex-C18 column (i.d. 100x2.1 mm, 1.7µm) with mobile phases comprising (A) water:acetonitrile:formic acid 63:37:0.02 (v/v/v) and (B) acetonitrile:isopropanol 1:1 (v/v).

### Statistical analysis

Statistical analysis was performed using GraphPad Prism 8.0. Mouse survival data between two groups were compared using Log-rank (Mantel–Cox) analysis. Mouse body weight change between two groups was compared using two-way ANOVA analysis. Other data from two groups were compared using the unpaired Student’s t test. P values less than 0.05 were considered statistically significant.

## Data Availability

The CRISPR screen and RNA sequencing data have been deposited in the National Center for Biotechnology Information Gene Expression Omnibus under accession code GSE298405, and are available from corresponding author upon reasonable request. All other study data are included in the article and/or supplementary materials.

## Acknowledgments

We thank Dr. Jing Han for kindly providing STING plasmid. This work was supported by Grants from the National Natural Science Foundation of China (82388201 and 32200751), the Chinese Academy of Medical Sciences Innovation Fund for Medical Sciences (2021-I2M-1-017, 2024-I2M-ZD-005) and the Young Elite Scientist Sponsorship Program by CAST (YESS20210066).

## Author Contributions

X.C. designed the research and supervised the study; S.L., J.W. performed the experiments. L.L., and J.Z. generated AAV9-Metabolic CRISPR library. S.L. and X.C. analyzed data and wrote the manuscript.

