## Supplementary materials for "*In vivo* AAV9-SB-CRISPR screen identifies fatty acid elongase ELOVL5 as a pro-resolving mediator in lung inflammation"

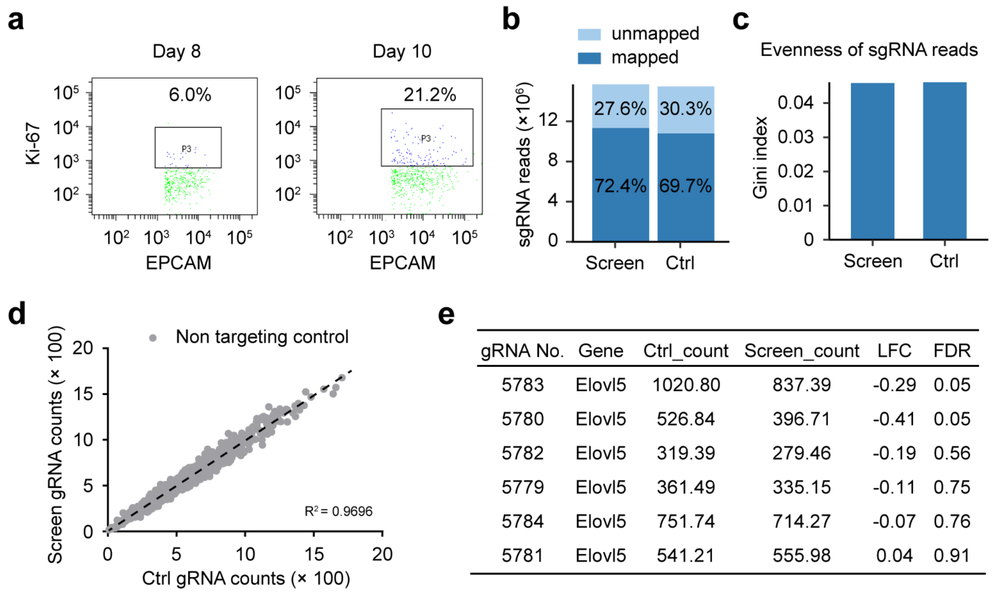


| gRNA No. | Gene | Ctrl_count | Screen_count | LFC | FDR |
|---|---|---|---|---|---|
| 5783 | Elovl5 | 1020.80 | 837.39 | -0.29 | 0.05 |
| 5780 | Elovl5 | 526.84 | 396.71 | -0.41 | 0.05 |
| 5782 | Elovl5 | 319.39 | 279.46 | -0.19 | 0.56 |
| 5779 | Elovl5 | 361.49 | 335.15 | -0.11 | 0.75 |
| 5784 | Elovl5 | 751.74 | 714.27 | -0.07 | 0.76 |
| 5781 | Elovl5 | 541.21 | 555.98 | 0.04 | 0.91 |

**Figure S1 *In vivo* CRISPR screen for essential metabolic genes in a mouse model of lung inflammation**

(**a**) Flow cytometry analysis of AECII proliferation in mice at different times post-IAV infection (50 PFU). (**b**) Mapping ratio of sequenced sgRNA reads in screen and control samples. (**c**) Evenness of sequenced sgRNA reads in screen and control samples. (**d**) Comparison of non-targeting control sgRNA read counts between resolving AECII (screen) versus resting AECII (control). (**e**) Read counts of *Elovl5* sgRNAs in screen and control samples.

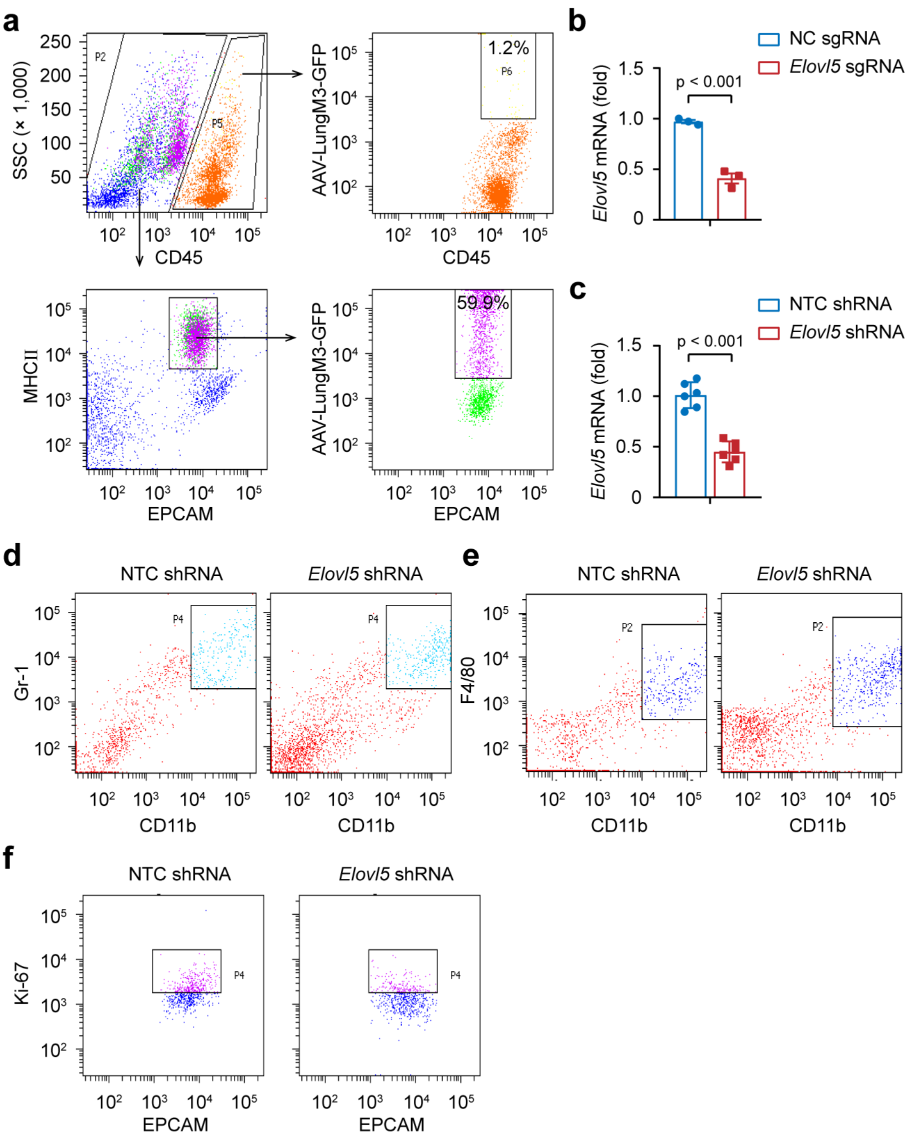


**Figure S2 ELOVL5 promotes lung inflammation resolution in mice after IAV infection**

(**a**) Flow cytometry analysis of infection efficiency of AAV-LungM3 in AECII and CD45$^+$ cells two weeks after transfection (5 × $10^{10}$ viral genome copies per mouse). (**b**) qPCR analysis of *Elovl5* mRNA expression in lungs of CAG-Cas9 mice after transfecting with AAV-LungM3 containing negative control sgRNA or sgRNA targeting *Elovl5* (5 × $10^{10}$ viral genome copies per mouse) for two weeks. (**c**) qPCR analysis of *Elovl5* mRNA expression in AECIIs from C57BL/6 mice after transfecting with AAV-LungM3 containing non-targeting control shRNA or shRNA targeting *Elovl5* (5 × $10^{10}$ viral genome copies per mouse) for two weeks. (**d**, **e**) Flow cytometry analysis of neutrophil (**d**) and macrophage (**e**) numbers in BALF from C57BL/6 mice at day 12 post-IAV infection (50 PFU) after transfecting with AAV-LungM3 containing non-targeting control shRNA or shRNA targeting *Elovl5* ($5 \times 10^{10}$ viral genome copies per mouse) for two weeks. (**f**) Flow cytometry analysis of proliferation of AECIIs from C57BL/6 mice described in **e**. Data are representative of three independent experiments (**a**-**f**), shown as mean ± SD in **b**, **c**, two-tailed unpaired Student's t test in **b**, **c**.

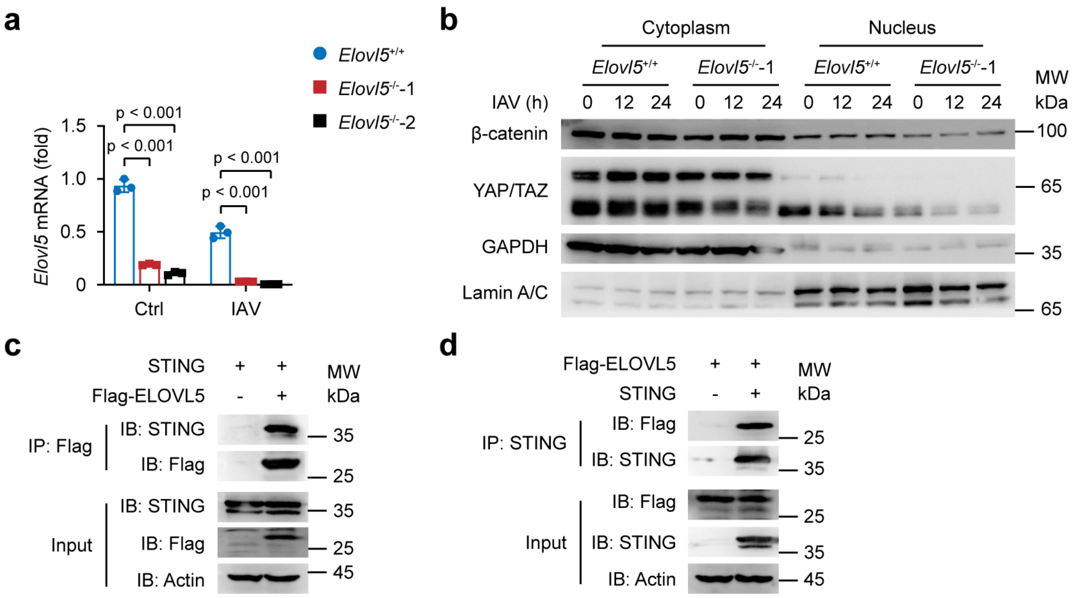


**Figure S3 ELOVL5 promotes inflammation resolution through interacting with STING in MLE-12 cells**

(**a**) qPCR analysis of *Elov5* mRNA expression in *Elovl5*$^{+/+}$ and *Elovl5*$^{-/-}$ MLE-12 cells infected with IAV (MOI = 1) for 24 h. (**b**) Immunoblot analysis of nuclear translocation of β-catenin and YAP/TAZ in *Elovl5*$^{+/+}$ and *Elovl5*$^{-/-}$ MLE-12 cells infected with IAV (MOI = 1) for the indicated times. (**c**, **d**) Immunoblot analysis of the interaction between ELOVL5 and STING in HEK293T cells transfected with Flag-ELOVL5 and STING-overexpressing plasmids. Data were representative of three independent experiments (**a**-**d**, shown as mean ± SD in **a**), unpaired Student's t test in **a**.

**Table S1. Primers used for q-PCR in this study.**

| Gene | Forward (5'-3') | Reverse (5'-3') |
|---|---|---|
| *Rpl32* | TTAAGCGAAACTGGCGGAAAC | TTGTTGCTCCCATAACCGATG |
| *Il10* | GCTCTTACTGACTGGCATGAG | CGCAGCTCTAGGAGCATGTG |
| *Tgfb2* | TCGACATGGATCAGTTTATGCG | CCCTGGTACTGTTGTAGATGGA |
| *Mki67* | GAGGAGAAACGCCAACCAAGAG | TTTGTCCTCGGTGGCGTTATCC |
| *Sftpc* | TCCTCGTTGTCGTGGTGATTG | GGAAAAGGTAGCGATGGTGTC |
| *Il1b* | CCCAACTGGTACATCAGCAC | TCTGCTCATTCACGAAAAGG |
| *Tnf* | AAGCCTGTAGCCCACGTCGTA | GGCACCACTAGTTGGTTGTCTTTG |
| *Ctgf* | GGGCCTCTTCTGCGATTTC | ATCCAGGCAAGTGCATTGGTA |
| *Cyr61* | CTGCGCTAAACAACTCAACGA | GCAGATCCCTTTCAGAGCGG |
| *Elovl5* | TTCGATGCGTCACTCAGTACCT | TGTCCAGGAGGAACCATCCTT |
